# Replicability of introgression under linked, polygenic selection

**DOI:** 10.1101/379578

**Authors:** Himani Sachdeva, Nicholas H. Barton

**Affiliations:** Institute of Science and Technology Austria (IST Austria), Am Campus 1, Klosterneuburg A-3400, Austria

**Keywords:** introgression, polygenic selection, linkage, parallel adaptation

## Abstract

We study how a block of genome with a large number of weakly selected loci introgresses under directional selection into a genetically homogeneous population. We derive exact expressions for the expected rate of growth of any fragment of the introduced block during the initial phase of introgression, and show that the growth rate of a single-locus variant is largely insensitive to its own additive effect, but depends instead on the combined effect of all loci within a characteristic linkage scale. The expected growth rate of a fragment is highly correlated with its long-term introgression probability in populations of moderate size, and can hence identify variants that are likely to introgress across replicate populations. We clarify how the introgression probability of an individual variant is determined by the interplay between hitchhiking with relatively large fragments during the early phase of introgression, and selection on fine scale variation within these, which at longer times results in differential introgression probabilities for beneficial and deleterious loci within successful fragments. By simulating individuals, we also investigate how introgression probabilities at individual loci depend on the variance of fitness effects, the net fitness of the introduced block, and the size of the recipient population, and how this shapes the net advance under selection. Our work suggests that even highly replicable substitutions may be associated with a range of selective effects, which makes it challenging to fine map the causal loci that underlie polygenic adaptation.

## Introduction

The extent to which phenotypic and genetic changes are replicated during adaptation across closely related populations has generated much interest (Conte et al. 2012; Storz 2016), and is part of the broader question of the predictability of evolutionary change (Lässig et al 2017). Parallel evolution can be investigated at multiple scales, and may refer to the involvement of the same mutations, the same nucleotides, the same genes, or even the same pathways during adaptive response in different populations (Manceau et al 2010). An important challenge is to interpret highly replicable genetic loci— do such loci necessarily make large contributions to the selected trait? Conversely, for different architectures of the selected trait, how often are the same genetic loci implicated across replicate populations?

Several factors can influence the replicability of allele frequency changes or adaptive substitutions at individual loci (Stern 2013). More shared variation between replicate populations makes it more likely that the same variants respond to selection across different replicates, but the response is then associated with a weaker reduction in neutral diversity, (i.e., ‘soft sweeps’, see Hermisson and Pennings 2005). By contrast, adaptive substitutions that arise de novo are rarely shared across replicates, but can be more easily identified as being under selection because of the associated hard sweep patterns.

Factors influencing genetic parallelism at the level of individual variants remain poorly understood for selection on highly polygenic traits, despite several studies with natural and laboratory replicates (Burke et al. 2010; Chan et al. 2012; Yeaman et al. 2016). Polygenic traits are often characterized by high genotypic redundancy, with multiple genotypes corresponding to approximately the same phenotype. For traits under stabilizing selection, a shift in the selection optimum can thus result in a highly heterogeneous response at the genotypic level, with selection amplifying initial random fluctuations in allele frequencies, to produce quite different outcomes across replicates, even if these initially share the same variation.

In general, adaptation in complex traits involves a large number of partially linked and weakly selected variants, and is thus characterized by pervasive hitchhiking– where effectively neutral or deleterious variants are swept to high frequencies along with clusters of positively selected variants (Barton 1995). While the effects of hitchhiking on neutral diversity have been studied extensively in the theoretical literature, most of these studies assume that selected variants are either all deleterious or all beneficial in a given population (e.g., Barton and Bengtsson 1986). However, a typical genome is likely to harbour many weakly selected and linked variants, both deleterious and beneficial. The efficacy of selection in discriminating between multiple, tightly linked beneficial and deleterious loci is an essential determinant of the total phenotypic response to selection in a finite population (Robertson 1970, 1977). It is also key to assessing the likelihood that an allele frequency change (even one that occurs across replicates) is itself adaptive and not a consequence of hitchhiking.

An important factor influencing the extent of hitchhiking (and the associated patterns of neutral diversity) is the density of selected polymorphisms on the genome. For example, Good et al. (2014) show that when the rate of deleterious mutations is higher than the typical selective effect per mutation, polymorphic variants are sufficiently common along the genome that neutral diversity no longer depends on the selective effects of individual variants. This results in a qualitatively different shape of the neutral site frequency spectrum, which cannot be predicted by standard models of background selection, but is better described by an *infinitesimal* framework, which is parametrized by the variance in population fitness (or more generally the variance per unit map length), rather than the variance of fitness effects at individual loci (Neher et al 2013).

In this paper, we focus on a similar scenario of a complex trait determined by a large number of weakly selected variants uniformly spread across the genome (see also Sachdeva and Barton 2018). The main goal is to explore how the interplay between linkage, polygenic selection and genetic drift shapes the long-term introgression probability of different fragments of such a genome, when it is introduced into a genetically homogeneous recipient population. For simplicity, we consider the introgression of a medium-sized block of map length *y*_0_ rather then the whole genome. This allows us to ignore certain complications, such as multiple crossovers within the selected region, in analytical calculations. It is also representative of a scenario where a genome that is repeatedly back-crossed into a recipient population, breaks up into several medium-sized segments, that evolve more or less independently, while rare in the recipient population. Most importantly, as we show below, the introgression probability of any variant depends primarily on the effect of variants within a characteristic linkage scale, and not on loosely linked regions, as long as the introduced genome has *no net* selective advantage in the recipient population. This again suggests that studying the introgression of medium-sized blocks can provide useful insight into the more general case.

The introduced block is assumed to carry a large number *L* of loci with a range of selective effects, uniformly spaced over map length *y*_0_. These loci contribute to an additive trait which is under directional selection in the recipient population. The trait value associated with any portion of the block is then just the sum of effects of all the loci it contains. The effect sizes of loci on the introduced block are drawn from a distribution with mean *μ* and variance *σ*^2^; the effect sizes of variants fixed in the recipient population can be set to zero without any loss of generality. For simplicity, we assume that *μ* and *σ*^2^ do not vary across the introduced block. However, most of our analytical results hold even when this condition is not met, for instance, if there is a statistically significant clustering of large-effect variants in a particular genomic window in the donor population.

If the donor population is in linkage equilibrium (LE), such that allelic states of different loci along the introduced block are statistically uncorrelated, then different segments of this block (each with *l*_1_ loci) have random (and typically unequal) contributions, with mean *μl*_1_ and variance *σ*^2^*l*_1_. Note that the variance of these contributions, or more generally, the variance per unit map length, given by *V*_0_=*σ*^2^*L*/*y*_0_, is the same irrespective of whether the block contains a few, large-effect loci (small *L*, large *σ*^2^) or many, small-effect loci (large *L*, small *σ*^2^). Thus, varying *L* while keeping *V*_0_ and *μL* constant, tunes the extent to which individual loci interfere with each other, while holding constant the average selective effect of fragments that span many such loci and have a given map length. If the number of loci is very large and the distribution of allelic effects correspondingly narrow, i.e., in the limit *μ*→0, *σ*^2^→0, *L*→∞, with *μL* and *σ*^2^*L* held fixed, this model approaches the *infinitesimal model with linkage* (see Robertson (1977), where it was first introduced). This model is similar to the well-known infinitesimal model (Bulmer 1980; Barton et al. 2017), but also accounts for linkage, by considering loci on a linear map, rather than unlinked loci.

In previous work, we analysed the initial introgression dynamics of a single block of genome introduced into a genetically homogeneous recipient population, under the infinitesimal model with linkage which assumes that the introduced block differs from ‘native’ blocks (belonging to the recipient population) at a very large number of weakly selected loci (Sachdeva and Barton 2018). This study showed that even if the introduced block has no overall selective advantage within the recipient population, the number of its descendant fragments and the total amount of introgressed genetic material, grows exponentially with time (assuming an infinite population). This is in spite of the fact that smaller and smaller descendant blocks are associated with weaker and weaker additive effects, and hence may be effectively neutral. Non-neutral (exponential) introgression dynamics arise nevertheless because of the emergence of medium-sized descendant sub-blocks, that are too small to undergo frequent recombination but large enough to have significant contributions to fitness and hence be amplified by selection.

In this study, we explore the effects that shape the introgression dynamics of different segments of a *particular* introduced block, in contrast to our previous work which was concerned with statistical averages associated with an *ensemble* of such blocks (all characterized by the same net trait value *z*_0_ and the same genic variance per unit map length *V*_0_). We focus here on the following two questions: first, what is the expected *initial* growth rate of a fragment or locus embedded within the introduced block, under directional selection? Is the growth rate of an individual variant influenced more by its own effect or the effects of linked loci within a characteristic map distance? Second, to what extent do expected initial growth rates predict the ultimate fixation probability of different fragments in a finite population?

The expected growth rates of genomic segments during the initial phase of introgression are relatively easy to calculate, since introgressed fragments are rare, and hence unlikely to encounter each other during mating. Thus, any genome carries at most one fragment of the introduced block, which leads to explicit analytical expressions for the initial growth rate of different fragments, and for the probability of survival of at least some part of the introduced block. We then simulate individuals to analyse long-term introgression in detail for one example, focusing on the correlation between initial growth rates and long-term fixation probability. Finally, we outline general trends, by considering how introgression probabilities at individual loci depend on the variance of fitness effects, the total trait value associated with the introduced block, the size of the recipient population, and the extent of initial stochasticity (which can be tuned in this model by introducing single versus multiple identical copies of the block). The extent to which selection and recombination can pick out and amplify individual favourable variants is a basic determinant of the net response to selection, when selected variants are tightly linked. We address the question of selection limits, and their dependence on population size and linkage, at the end of the paper.

The distinction between initial and long-term introgression is related to the qualitatively different role played by recombination during these two phases (Sachdeva and Barton 2018). While recombination primarily breaks apart the introduced genome into smaller fragments during the initial phase of introgression, over longer times, as these fragments become more abundant in the population, recombination can also bring together various successful fragments, countering Hill-Robertson interference. Thus, over these time scales, recombination can uncover fine scale variation within successful fragments (provided these have not fixed), allowing selection to discriminate between tightly linked variants. In a very large population, this would allow selection to fix all positively selected variants and eliminate all deleterious variants within introgressing fragments that have survived the initial phase. In a smaller population, only part of this fine-scale discrimination can be achieved, which in turn constrains the net response to selection. An important focus of our work is to explore how the ultimate fate of individual variants on the genome in a finite population is determined by the interplay between the early phase dynamics characterized by selection on relatively large fragments of the genome and long-term dynamics governed by selection on fine scale variation within these. The broader aim is to clarify the conditions under which individual variants have high establishment probability (and hence a strong likelihood of being replicated across populations which receive the same genome). Identifying the distribution of effects associated with highly replicable loci, even within this simple model, can shed light on the possibilities and limitations of fine mapping the causal loci that underlie polygenic adaptation.

For simplicity, we ignore several factors that can qualitatively influence replicability, such as epistatic interactions between loci and de novo mutation. We also assume that all replicate populations receive *identical* genomes from the donor population; this results in a much higher degree of convergence than would occur across natural replicates, where introgressing genomes would share only some (but typically not all) variants. Further, multiple genomes introduced into a single population are also assumed to be all identical. In general, heterogeneity among introduced genomes from the donor population or within the recipient population would result in much lower establishment probabilities of individual variants, and thus less convergence. We comment on possible extensions of the model that relax some of these assumptions in the Discussion.

## Model and Methods

Consider a situation where *N*_0_ *identical* copies of a block of genome are introduced into a very large population of diploids at *t*=0. Any diploid individual is assumed to carry at most one copy of the introduced block. The block has *L* loci which are assumed to be evenly spaced on the genetic map, with recombination rate *c* between adjacent loci. The extension to unequal rates of recombination between loci is straightforward. The additive effect of the *k^th^* locus is denoted by *s_k_*. The fitness of the block is the sum of effects of all the loci, 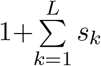.

### Initial introgression into a large population

To derive analytical results for the initial dynamics of block fragments, we assume that the map length of the introduced block, given by *y*_0_=*cL*, is small enough that multiple crossovers can be neglected. If we further assume that the introduced block and its descendant fragments are sufficiently rare in the population (as expected during the initial phase), then the probability of recombination between fragments is negligible, and any individual inherits at most one introgressed fragment. Therefore, we need only consider single fragments spanning loci *i* ⋯ *j*, with fitness 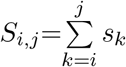, and which break up by recombination at rate *c*(*j*−*i*). If selection and recombination are weak, the expected numbers *N_i,j_*(*t*) of different fragments change approximately continuously through time, according to a set of linear equations:

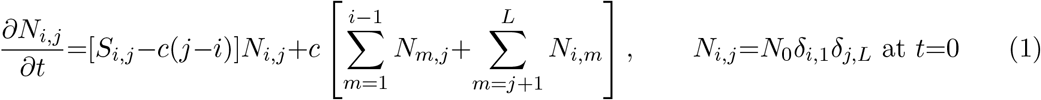

The first term is the rate at which a fragment spreads intact (without being split), and is positive when the fragment is amplified by selection (at a rate proportional to its fitness *S_i,j_*) faster than it is split by recombination (at a rate proportional to map length *c*(*j*−*i*)). The second and third terms are due to generation of the fragment by recombination from larger blocks (in which it is embedded).

Equation (1) can be solved explicitly by taking the Laplace Transform, solving for the expected number of single recombinants *N*_1, *i*_ and *N_i,L_*, and then solving for smaller blocks. The solution is a sum of exponentially decaying (or increasing) components:

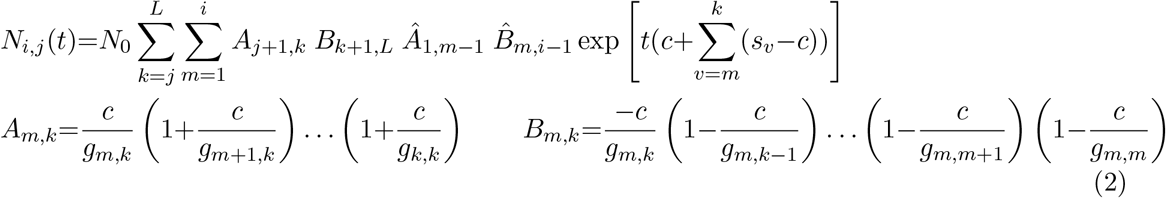

where 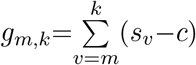; *Â* is obtained by replacing *c* by −*c* in *A*, and 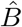 by replacing −*c* by +*c* in *B*. Also, *A_m,k_*=1 for all *m*>*k*, and so on for *B_m,k_, Â_m,k_*, 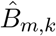.

The sum is over all blocks [*m, k*] (with *m*≤*i*≤*j*≤*k*) that contain the fragment [*i,j*], including [*i, j*] itself. For large *t*, the exponential term with the largest growth rate will dominate the dynamics of the block. Thus, at long times, any block grows at the same rate as the fastest growing parent block from which it can be generated. Note that if the intrinsic growth rate *S_i,j_*−*c*(*j*−*i*) of the block is larger than the growth rate of any parent block containing it, then the long term growth rate of the block is just *S_i,j_*−*c*(*j*−*i*). An important consequence of eq. (2) is that deleterious fragments may also spread through the population, if generated at a constant rate by a beneficial, exponentially growing parent block. In fact in this case, the deleterious sub-block will grow at the same rate as the beneficial parent block, as evident from eq. (2). Thus, each such exponentially increasing block generates various descendant sub-blocks of varying fitness, giving rise to a family of blocks, akin to a quasi-species (Eigen et al. 1988).

Figure 1A depicts the long term growth rates of different fragments of a block with 10 loci. Figure 1B shows the expected numbers of all possible 55 fragments contained within this block as a function of time— black lines represent fragments with a negative long-term rate of change, while coloured lines represent fragments with positive long-term growth rates. All fragments with the same long-term (positive) growth rate are represented by one colour. Figure 1B thus suggests that the introgression of a block of genome under directional selection results in the spread of different families of sub-blocks at different rates. Each such family (represented by lines of one color) consists of sub-blocks that are all contained within a particular, fast growing parent block. After an initial transient phase, all sub-blocks within a family increase at the rate of growth of this parent block, at least within a very large population in which introgressing blocks do not encounter each other. Note that the growth rates discussed here describe the dynamics of the *expected* number of copies of any fragment in the population, averaging over all possible stochastic histories, including those in which the fragment is lost.

**Figure 1:**
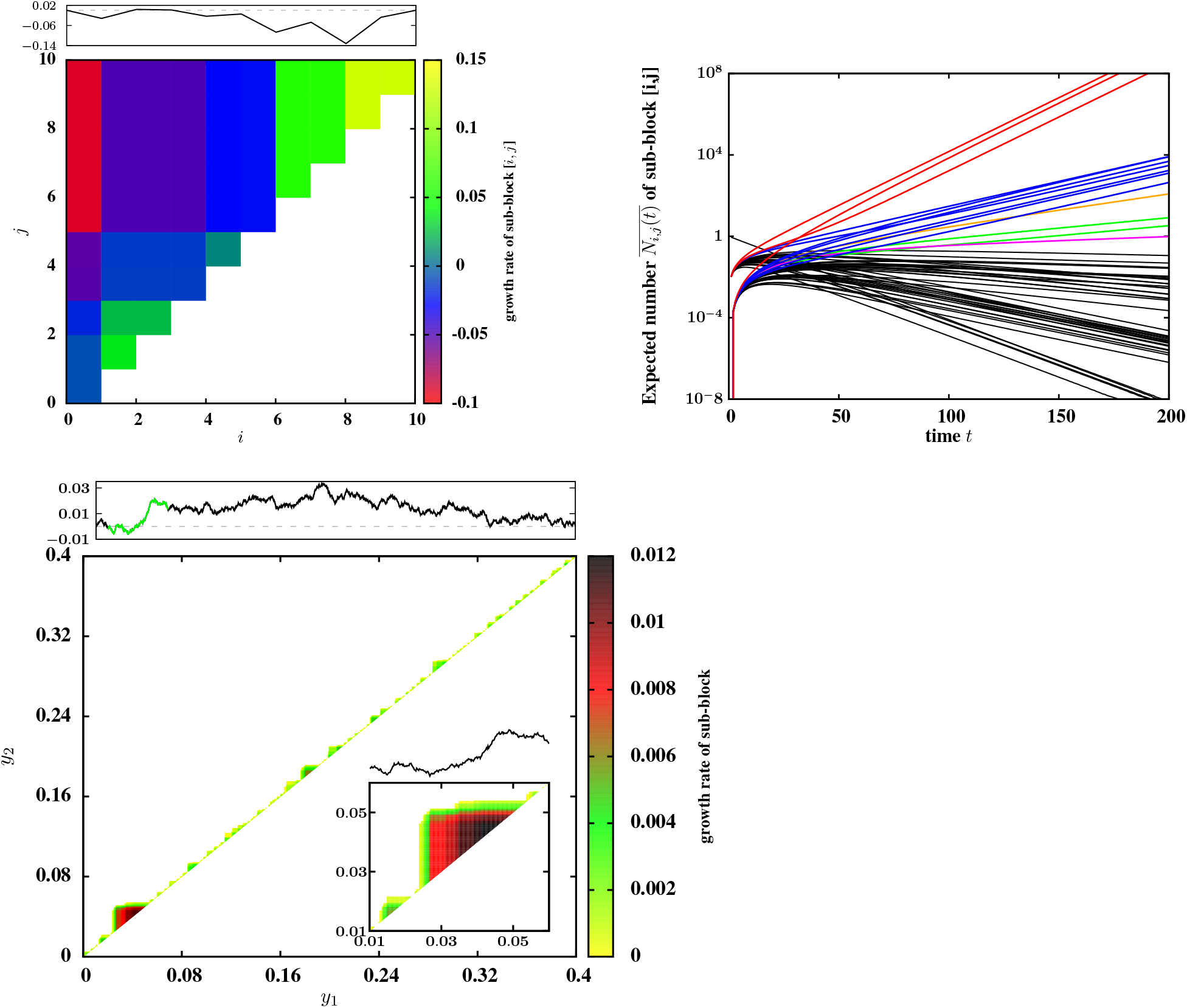
(A) Expected (asymptotic) growth rates of various fragments [*i,j*] of a block with *L*=10 loci, with effect sizes drawn from a Gaussian distribution with mean *μ*=0 and variance *σ*^2^=0.1/*L*, subject to the constraint that the net effect of the block is 0; the rate of recombination between adjacent loci is *c*=0.01. The two axes of the plot represent the two end-points of the fragment; each location (*i, j*) in the upper triangle of the plot represents a particular fragment with end points *i* and *j*≥*i*. the colour associated with each location encodes the expected growth rate of the fragment (see adjoining color scale). Note that many of the 55 sub-blocks have the same expected asymptotic growth rate; thus there are 15 distinct asymptotic growth rates (colors) in (A). The black curve in the upper panel shows how the trait value varies along the introduced block. The x-axis represents the positions *i* of the loci on the block; the y-axis represents the cumulative contribution of all loci from [0, *i*]; thus any segment with a positive slope represents a sub-block with a net positive contribution to trait value and vice versa. (B) Expected numbers of each of the 55 fragments contained within the 10-locus block in (A), as a function of time *t*. Black curves represent fragments with asymptotically negative growth rates; curves in other colors represent fragments with asymptotically positive growth rates. All fragments with the same asymptotic (positive) growth rate are represented by the same color. All curves are calculated by iterating eq. (1). (C) Asymptotic growth rates of various fragments of a block with *L*=2048 loci and map length *y*_0_=0.4. The net trait value associated with the block is 0 and variance per unit map length 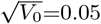 (which implies *σ*^2^=0.001/*L*). The two axes of the plot represent the map positions *y*_1_ and *y*_2_ of the two end-points of any fragment {*y*_1_, *y*_2_} in the block. Only those fragments that have a positive expected growth rate in the long run are shown. The inset inside the main plot zooms into a particular region of the block (highlighted in green in the upper panel) and depicts fragments that have a positive growth rate within this region.

In the following, we will approximate the growth rate of any fragment by the rate associated with the fastest growing term in eq. (2), i.e., by the intrinsic growth rate of the fastest growing parent block that contains this fragment. We refer to this as the ‘expected’ growth rate of the fragment.

The key assumption underlying eq. (1) is that descendants of the introduced fragment always form a negligible fraction of the population, and thus never encounter each other during mating. Then the spread of introgressed genetic material through the population can be formulated as a *branching process* (BP) that treats the (forward in time) lineages of different descendants as being independent of each other, but having a common dependence on (*s*_1_, *s*_2_,… *s_L_*} (Sachdeva and Barton 2018). The BP framework is very powerful and can, in principle, be used to calculate the full distribution of introgressing fragments as a function of time. Here, we write down the equation for the probability *P*_1, *L*_ that at least *some* part of a block {1, *L*} spanning loci *i*=1, ⋯ *L* survives at long times, given that it is initially introduced as a single copy. It is convenient to work with dimensionless quantities, and so we scale both selection and survival probability relative to the map length 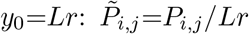, and 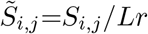. The long-term scaled survival probability of a block (1, *L*} satisfies:

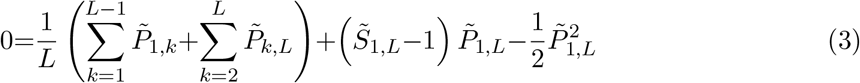

This equation is of the form 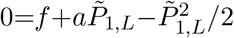, where the first term *f* is due to the survival of smaller sub-blocks, and the coefficient *a* of the second term is the intrinsic growth rate of the full block. This equation is analogous to eq. 3 in Barton (1995), but has an additional driving term *f*. Solving the quadratic, we have:

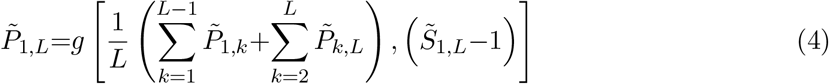

where 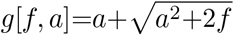; *g*[0, *a*]=*max*[0, 2*a*]. This is easily evaluated, because 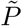 for larger blocks depends only on that for smaller blocks. Note that blocks with a negative net rate of increase (*a*<0) can survive (in part), if they contain smaller blocks that have a positive rate of growth, such that *f*>0. If the block {1, *L*} is introduced in *N*_0_ copies, then we can treat the survival probability of each such block as independent of other blocks, as long as *N*_0_ is much smaller than the size of the recipient population. Then the overall survival probability is just 1−(1−*P*_1, *L*_)^*N*_0_^.

### Long-term introgression

Equation (1) is a valid description of the dynamics of individual fragments only while any individual carries at most one such fragment. As shown in Sachdeva and Barton (2018), this is true over a time scale that scales weakly (~log(*N*)) with the size *N* of the recipient population. Beyond this initial time scale, mating between individuals carrying introgressed material becomes more frequent and genomes carrying multiple fragments of the introduced block emerge. This phase is thus characterized by competition between genomes bearing different mosaics of the fragments that have established and proliferated in the initial phase, and is no longer described by eq. (1).

To study long-term introgression, we simulate populations with *N* diploid individuals. The simulations are initialized by assuming that 2*p*_0_*N* haploid genomes in the population carry identical ‘introduced’ blocks, while all other genomes carry identical ‘native’ blocks. For simplicity, we assume that each of the 2*p*_0_*N* introduced blocks is present in a different individual. Each block has *L* loci, with rate of recombination *c* between adjacent loci. The allelic effect of each native locus is equal to zero, while effects of the introduced loci are drawn from a distribution with variance *σ*^2^ and mean *μ*. The trait value *S* associated with any individual is then just the sum of effects of all introduced variants in its diploid genome; the fitness of the individual is *e^S^*.

In each generation, 2*N* parents are drawn by randomly sampling individuals in proportion to their fitness. Each parent produces a gamete with recombination between parental haplotypes— the number of crossover points is drawn from a Poisson distribution with mean *y*_0_; the locations of the crossover points are chosen by uniformly sampling one of the *L*−1 junctions between loci without replacement. The gametes are then paired to form *N* individuals of the next generation.

We compute introgression probabilities by performing 200–400 replicate simulations, each initialized by introducing 2*p*_0_*N* identical copies of the same block at *t*=0. Thus, replicate populations differ only in the stochastic history of reproduction and recombination. The introgression probability of a particular fragment at time *t* is calculated as the fraction of replicates in which the fragment is present *either by itself or as part of a larger block*. Since any fragment must either fix or be completely eliminated at long times, the introgression probability of a fragment in the lathe *t* limit, is just its fixation probability. By definition, any fragment with high introgression or fixation probability is also a ‘replicable’ fragment— a fragment with fixation probability *P* would be found in two replicate populations with probability *P*^2^. Note that the *introgression probability* is not the same as the *survival probability P*_1, *L*_, which was calculated in eq. (4). The latter refers to the probability that the block {1, *L*}, introduced as a single copy at *t*=0, is not fully lost from the population at long times, but has at least one surviving descendant sub-block.

### Data availability statement

FORTRAN 95 codes used to generate the simulated data can be found at: https://git.ist.ac.at/himani.sachdeva/source_codes_replicability_introgression_patterns/snippets.

## Results

To what extent is the ultimate fixation probability of different parts of the introduced genome shaped by the early phase dynamics (as encapsulated by the expected initial growth rates), and how is it affected by later phase dynamics (which are characterized by selection on multiple, linked ‘successful’ fragments)? To gain some insight into these issues, we first analyse one example in detail.

### Introgression probabilities of different fragments of the genome: an example

Consider an introduced block with *L*=2048 loci, uniformly spaced over map length *y*_0_=0.4. The contributions of different loci are drawn from a normal distribution with mean 0 and variance *σ*^2^=0.001/*L*, using an iterative scheme that ensures that the contributions sum to *S*_1, *L*_=0 (see also Sachdeva and Barton 2018). Figure 1C depicts all fragments of this block that have a positive expected growth rate. Note that these fragments are much smaller than the full block. By contrast, longer fragments of a nearly neutral introduced block are split by recombination faster than they are amplified by selection, and thus have negative expected growth rates.

In this example, we simulate a diploid population of size *N*=4000, with initial frequency of the introduced haplotype at *p*_0_=0.005, such that there are 2*Np*_0_=40 introduced blocks in the population at *t*=0. This corresponds to having 2*p*_0_*N* approximately independent realizations of the initial introgression process within any one population. Thus, the dynamics of early introgression in a population is expected to be close to the prediction of eq. (2) for 2*p*_0_*N*≫1. By contrast, if just one block is introduced (*p*_0_=1/2*N*), then all or some of the introduced genome, including beneficial fragments, may be lost from the population, while present in low numbers, which causes the dynamics of individual populations to differ markedly from the expected dynamics. The parameter *p*_0_ thus governs the extent to which the initial dynamics of any individual population is deterministic.

Figures 2A-2B show the introgression probability of different fragments of the introduced genome versus their expected growth rates (as calculated above, see eq. (2) and subsequent discussion) for two different time instants: *t*=600 (fig. 2A) and *t*=24000 (fig. 2B). Each point represents a particular fragment; the color of the point encodes the length of the fragment or alternatively, the number of equally-spaced selected loci that it contains (see accompanying color scale). The introgression probability of any fragment is the fraction of replicates in which the fragment is present, considering only those replicates in which at least some part of the introduced block survives at long times. Thus, introgression probabilities shown in fig. 2A and 2B are normalized with respect to the survival probability *P*_1, *L*_(*t*) (i.e., the probability that the introduced block is not fully lost from the population by time *t*). We consider here only introgressing fragments with significant average frequency (greater than 0.1), as these are unlikely to be lost by stochastic fluctuation.

**Figure 2:**
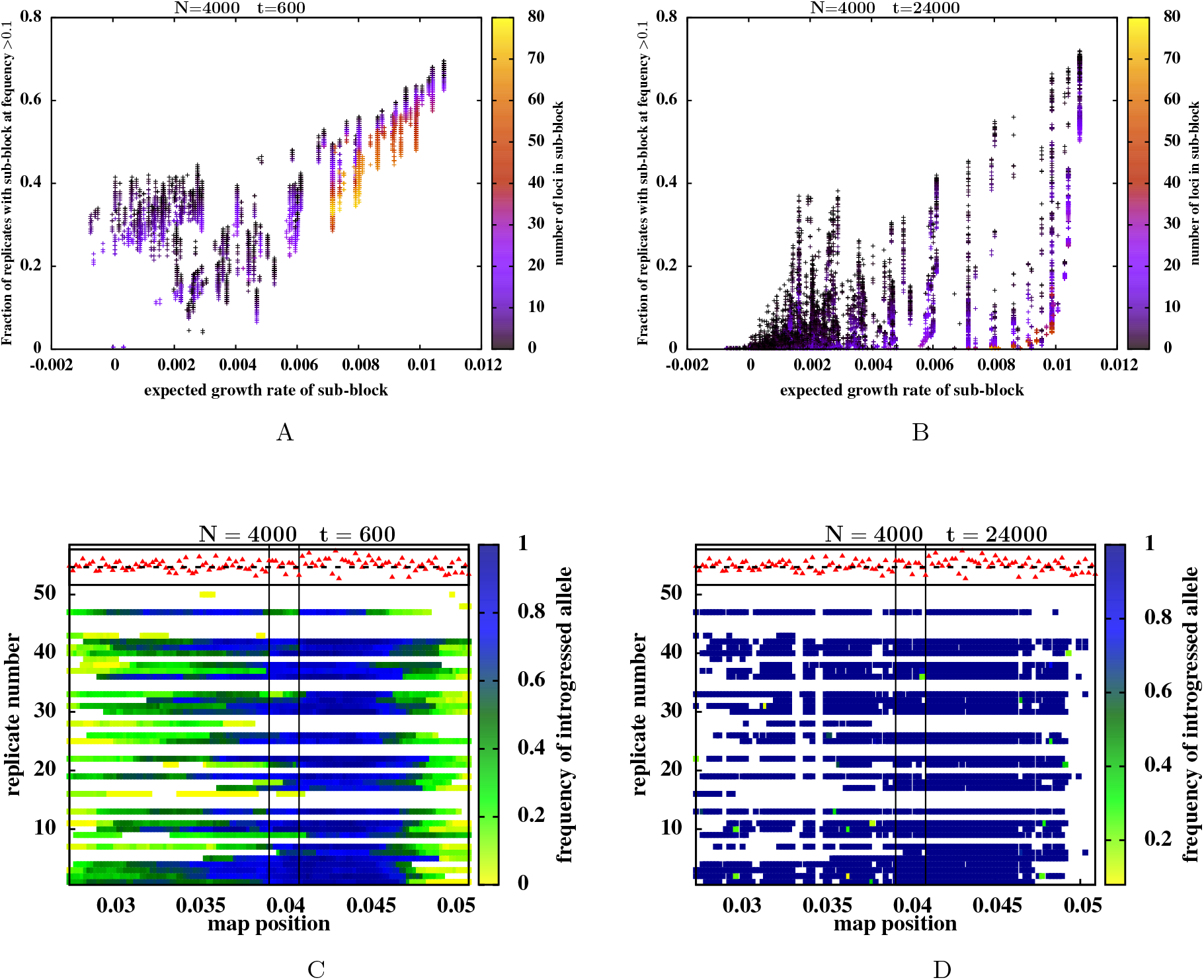
Introgression probabilities of different *genomic fragments* in a finite population. (A)-(B) The probability that a sub-block is present (either by itself or as part of a larger block) at frequency greater than 0.1 in a replicate, at *t*=600 (fig. A) and at *t*=24000 (fig. B), versus the expected growth rate of the sub-block. Each point represents a sub-block; the color represents the length of the sub-block (or alternatively the number of equally-spaced loci it contains). Note that at *t*=24000, sub-blocks with the same expected growth rate can have significantly different introgression probabilities. These are negatively correlated with the length of the sub-block. (C)-(D) Snapshots of 50 replicates carrying at least some introgressed material at *t*=24000 (fig. D) and the same replicates at *t*=600 (fig. C). The x-axis denotes the map position along the genome (for a limited genomic stretch); each horizontal line depicts a different replicate population (numbered 1 to 50 along the y-axis); the colored segments within each horizontal line represent fragments of the introduced block present in the population at time *t*; the color represents the frequency of the introgressed variant at that genomic position in that population. All replicate populations receive the same introduced genome at *t*=0; the selective effect of loci on this genome are depicted by the red triangles. All triangles lying below *s*=0 (the dashed black line) are deleterious. The window between the two vertical lines represents the sub-block with the second highest expected growth rate (among all sub-blocks contained in the introduced genome). All plots are obtained from individual-based simulations of a population of size *N*=4000; different replicate populations receive the same introduced block (with *S*_1, *L*_=0, *y*_0_=0.4, 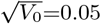, *L*=2048) at *t*=0, at frequency *p*_0_=0.005. Expected growth rates are calculated as described in the text.

The introgression probabilities of different fragments at *t*=600 appear to be highly correlated with their expected growth rates (correlation coefficient *r*=0.808), while the correlation is somewhat weaker (*r*=0.447) but still non-zero at *t* = 24000. The weaker correlation at longer times is due to significant differences in the long-term introgression probabilities of different fragments which have the *same* expected growth rate (note the much higher scatter along the y-axis in fig. 2B relative to fig. 2A).

Figure 2B suggests that the introgression of shorter fragments is more replicable than that of longer fragments, within any family of fragments with the same expected growth rate: note the higher introgression probabilities associated with black as compared to blue points within each vertical column of points. To investigate this in more detail, we zoom into a small window of the genome that contains a large number of replicable fragments. Figures 2C and 2D show snapshots of this genomic window for 50 replicate populations at *t*=600 and *t*=24000 respectively. Each horizontal band corresponds to one replicate population, the colors along each such band encode the frequencies (in that population) of introgressed variants at different map positions. The red triangles depict selective effects of individual variants.

At long times (fig. 2D corresponding to *t*=24000), *multiple*, disjoint fragments of the introduced genome are fixed in any population, even within this limited genomic region. We focus on the genomic window between the two vertical lines: this segment has the second highest expected growth rate among all the segments of the introduced genome. Note that many of the replicate populations lack a single, deleterious introgressed variant in the middle of this segment. These populations most probably witnessed one (or a few) recombination events that brought together two fragments of this segment onto a single genome *without the deleterious variant* in the middle. This new combination of fragments would then have out-competed the original segment, as it is associated with a larger selective advantage than the original segment, while having the same (low) probability of being split by recombination. In a subsequent section, we analyse this process in more detail for a toy example with three loci.

To avoid comparing fragments of different sizes, many of which are overlapping or even fully contained within other fragments, we compare introgression probabilities of different *single-locus variants* embedded within the introduced genome. Figures 3A and 3B show introgression probabilities at single loci (which constitute a subset of the fragments shown in figs. 2A and 2B) versus their expected growth rates, for *t*=600 and *t*=24000. As before, the expected growth rate of a locus is the growth rate of the fastest-growing sub-block containing it. The introgression probability at individual loci shows high correlation with the expected growth rate, at both *t*=600 (with correlation coefficient *r*=0.752) and at *t*=24000 (with *r*=0.819). At longer times, however, there is a significant scatter among loci with the same expected growth rate, with deleterious loci (purple dots) having lower introgression probabilities than beneficial loci (orange dots) within any particular cluster of such loci.

**Figure 3:**
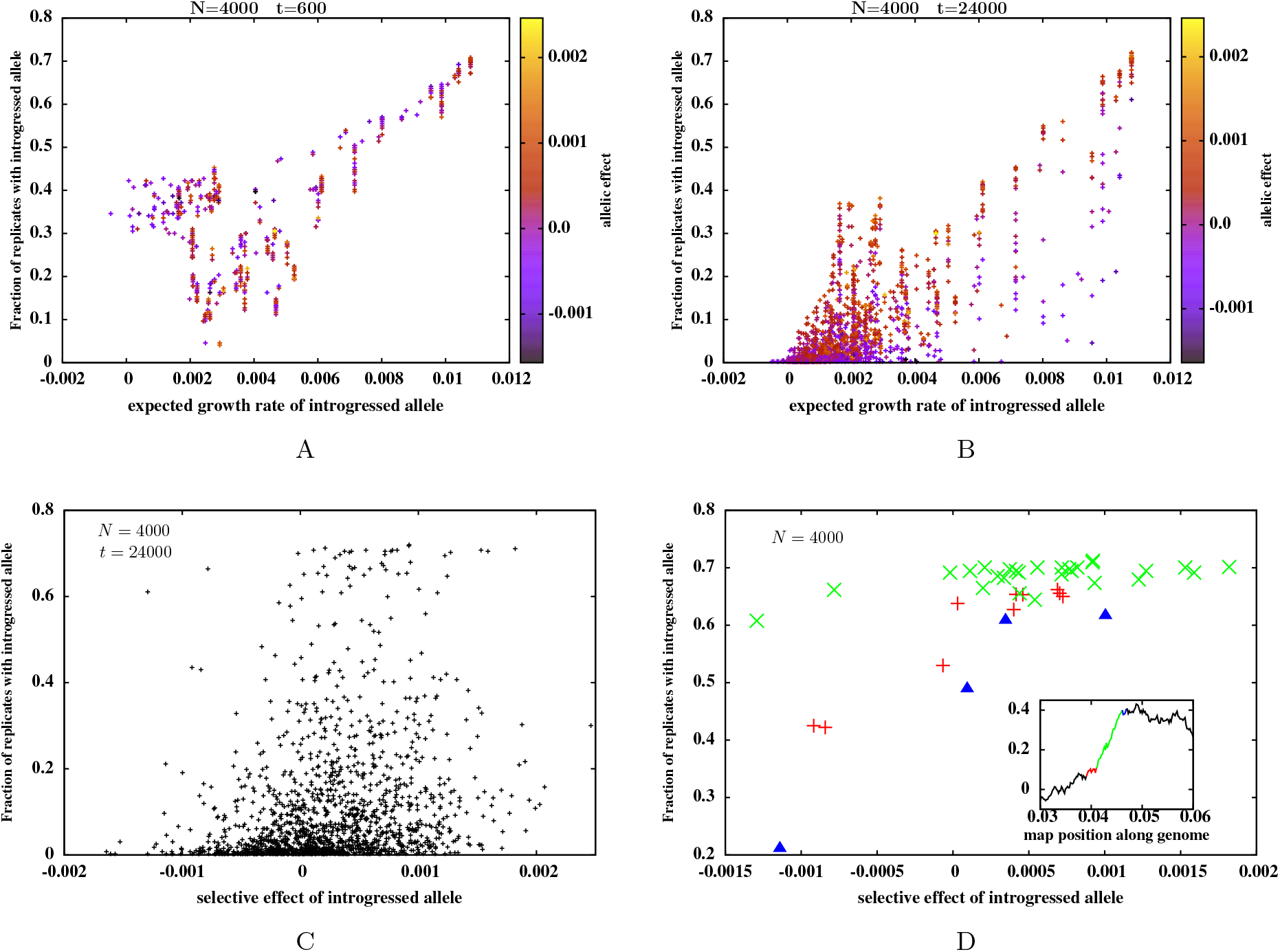
Introgression probabilities of different *single-locus variants*. (A)-(B) The probability that a particular single-locus variant (that was originally part of the introduced block) is present either by itself or as part of a larger block, at frequency greater than 0.1 in a recipient population, at *t*=600 (fig. A) and at *t*=24000 (fig. B), versus its expected growth rate. Each point represents a locus; the color represents the selective effect of the introduced variant at this locus. Note that at *t*=24000, loci with the same growth rate can have significantly different introgression probabilities, which appear to be negatively correlated with the selective effect of the loci. (C)-(D) The probability that a particular introduced, single-locus variant is present (either by itself or as part of a larger block) at frequency greater than 0.1 in a recipient population, at *t*=24000, versus the selective effect associated with the locus for: (C) all single-locus variants contained in the introduced block, and (D) single-locus variants present in three windows or regions within the introduced block. These three genomic windows are represented in the inset of fig. D. The inset shows the variation of trait value along a part of the introduced block, with the green, red and blue segments depicting sub-blocks which are associated with the three highest expected growth rates— 0.01078, 0.01041, and 0.01029 respectively. The green, red and blue points in the main plot represent introgression probabilities of single-locus variants contained in these three segments. Note that the introgression probability is very weakly correlated with the selective effect across all loci (fig. C), but displays a much stronger correlation within clusters of loci that have the same expected growth rate (fig. D). All plots are obtained from individual-based simulations of a population of size *N*=4000 by averaging over replicate populations receiving the same introduced block (*S*_1, *L*_=0, *y*_0_=0.4, 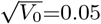, *L*=2048) at *t*=0 at frequency *p*_0_=0.005. All parameters are as in fig. 2.

To explore how long-term introgression probabilities at individual loci depend on their own selective effects, we consider three separate windows of the introduced genome (shown in green, red and blue in the inset of fig. 3D) These three segments depict the three fragments of the introduced genome that are associated with the three highest expected growth rates (see caption of fig. 3 for more details). The main plot in fig. 3D shows the long-term introgression probabilities of single-locus variants contained within each of these three segments, versus their selective effects, with green, red and blue points depicting loci contained in the corresponding segments shown in the inset. Figure 3D suggests that among loci with the same expected growth rate, long-term introgression probabilities are sensitive to the selective effect of the locus. In particular, the introgression probabilities of deleterious loci appear to decline in proportion to their selective disadvantage (note especially the red and blue points).

In the early phases of introgression, tightly linked loci, i.e., loci contained within the same exponentially increasing sub-block, have the same fate (as evident in the limited scatter of introgression probabilities among loci with the same expected growth rate in fig. 3A). At early times, selection does not ‘see’ individual loci, since recombination has not had sufficient time to separate tightly linked variants from each other. However, at longer times, recombination generates smaller and smaller fragments of successful sub-blocks and also reconstruct various combinations of these fragments, some of which would lack deleterious variants present in the original parent block. Such combinations would supplant the original successful block, thus also eliminating some of the deleterious loci contained within this block. Note that this kind of fine-grained separation of tightly linked beneficial and deleterious variants within a block over long time scales can only occur if the block has not already fixed in the population. This implies that the higher the expected growth rate of a sub-block, the sooner it would fix, resulting in much less variation between the introgression probabilities of its constituent loci. This is indeed what we see in fig. 3D, where the introgression probabilities of loci within the fastest growing sub-block (green points) depend weakly on selective effect, while loci on sub-blocks with the second and third highest expected growth rates (red and blue points respectively) exhibit a much stronger dependence.

Does the significant correlation between introgression probabilities and selective effect at individual loci within *limited* genomic windows, lead to a correspondingly high correlation across larger map distances? Figure 3C shows introgression probabilities (at *t*=24000) of single-locus variants across the entire block, versus their selective effects. Even in the long run, single-locus introgression probabilities are only weakly correlated with selective effect across the whole block (correlation coefficient *r*=0.246 in fig. 3C). Contrast this with the much stronger correlation with expected growth rates (*r*=0.819 in fig. 3B). This is consistent with our explanation that the ultimate fixation probability is shaped by linked selection (or selection on clusters of loci) in the early phases of introgression, which then constrains the extent to which selection can distinguish between individual variants within these clusters in the later phases of introgression. Note further that the most replicable variants are associated with a range of selective effects (see also fig. 4), and can even be significantly deleterious, which makes it misleading, at least within this model, to infer that high replicability implies adaptive significance.

**Figure 4:**
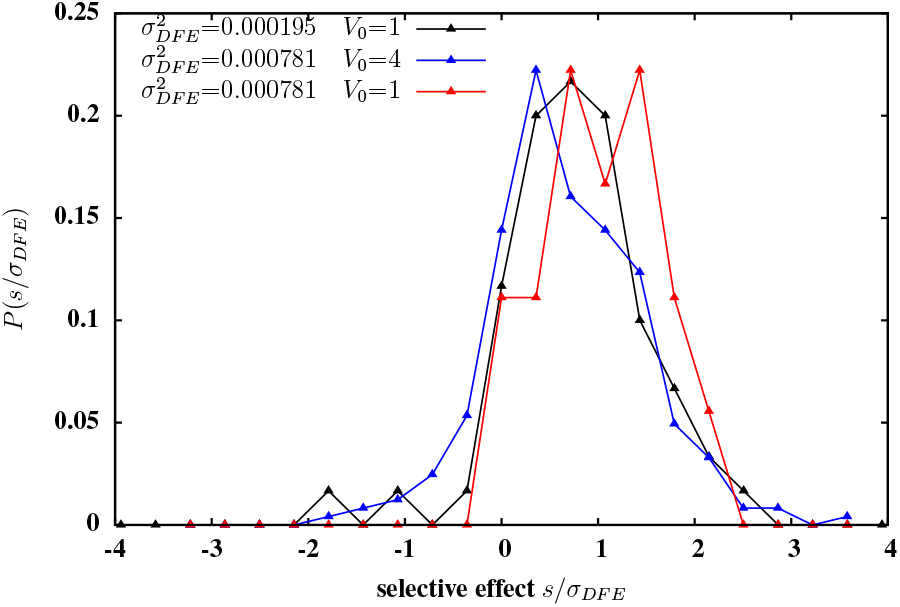
The distribution of selective effects among loci with greater than 50% probability of introgression. The three distributions correspond to three different introduced blocks with map length *y*_0_=0.4 and *S*_1, *L*_=0.0. The first block (distribution in black) has *L*=2048 individual loci with allelic effects drawn from a distribution with variance 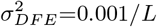, resulting in genic variance per unit map length 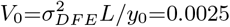. The second block (blue) has *L*=2048 loci, with the allelic effect of each locus exactly twice the effect of the corresponding locus in block 1; thus block 2 has 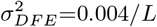 and *V*_0_=0.01. The third block (red) has L=512 loci; the allelic effect of the *i^th^* locus in this block is 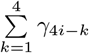, where *γ_i_* is the effect of the *i^th^* locus on block 1; thus block 2 has 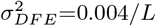 but *V*_0_=0.0025. Note that loci with negative selective effects are least likely to establish for the third block (red) that has fewer and larger effect loci. All plots are obtained from individual-based simulations of a population of size *N*=4000; the introduced block enters the population at *t*=0 at frequency *p*_0_=0.005.

### Dependence of introgression probabilities on the distribution of fitness effects

This raises the question: are long-term introgression probabilities of single locus variants more strongly correlated with their own selective effects, when these effects are larger? In other words, as the distribution of fitness effects becomes wider, is there an approach to an effective single-locus regime, where selection on the focal locus, rather than linked selection, determines introgression outcomes? In examining this question, we must distinguish between the variance of fitness effects *σ*^2^, and the closely related genic variance per unit map length *V*_0_=*σ*^2^/*c*, which depends on both *σ*^2^ and the density of selected loci on the genome. The genic variance *V*_o_ influences the extent of linked selection— a fragment of length *y* (emerging from an introduced genome with *S*_1, *L*_~0) has a typical (positive or negative) contribution that scales as 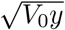. Thus the typical expected growth rates associated with successful fragments of the introduced genome are higher, and the time to fixation of these fragments correspondingly shorter, when *V*_0_ is larger. On the other hand, the per locus variance *σ*^2^ determines the extent to which selection can discriminate between loci of different effects at fine recombination scales at long times. Thus, large *σ*^2^ and large *V*_0_ are expected to influence the correlation between introgression probabilities and selective effect differently— while larger *σ*^2^ implies higher efficacy of selection in separating tightly linked beneficial and deleterious loci, higher *V*_0_ results in higher growth rates of successful blocks, which causes tightly linked loci to fix together before they can be taken apart by recombination and seen individually by selection.

To clarify the roles of *σ*^2^ and *V*_0_ in determining long-term introgression probabilities, we compare two blocks with the same density of selected variants (i.e., with the same number *L* of selected loci, and the same recombination rate *c* per locus), but with the variance of fitness effects of individual loci on one block being four times the corresponding variance for the other block. It then follows that the genic variance per unit map length *V*_0_ for the first block is also four times the corresponding variance for the second block. We consider all variants with longterm introgression probabilities greater than 0.5 within each block, and plot the distribution of their selective effects (fig. 4). Note that the distribution has similar shape in the two cases, but is shifted towards slightly *weaker* selective effects (relative to *σ_DFE_*) when the block has a wider DFE and a correspondingly high genic variance *V*_0_ (blue curve), than when the block has a more narrow DFE and a lower value of *V*_0_ (black). The mean selective effect of such significantly replicable loci is 0.662*σ_DFE_* in the first case (blue curve) and 0.754*σ_DFE_* in the second case (black curve). Moreover, the fraction of significantly replicable loci that are *deleterious* is slightly higher (0.156 versus 0.117), for the introduced block with larger 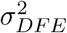 and *V*_0_. This suggests that for a given (high) density of selected variants on the genome, a wider DFE results in more extensive hitchhiking of deleterious loci during introgression, and thus reduced power to pinpoint adaptive loci from patterns of replicability.

We also compare two blocks characterized by the same *V*_0_, but with different values of *σ*^2^_*DFE*_ and *L*. Since *V*_0_ governs initial introgression, both blocks are expected to show very similar short-term patterns of introgression. However, a population that receives the block with wider DFE (and lower density of selected variants) should undergo more efficient elimination of deleterious loci in the long run. This is indeed what we see in simulations— the distribution of selective effects among significantly replicable loci on a block with wider DFE (red curve) is shifted towards higher effects (mean selective effect 1.0285*σ_DFE_*), relative to the corresponding distribution for the block that has a more narrow DFE but the same *V*_0_ (black curve). Further, the fraction of significantly replicable loci that are *deleterious* is negligible for the red curve, as compared to 0.117 for the black curve. This is consistent with the general expectation that when the additive contribution of a genomic region is concentrated at fewer, larger effect loci, then selection acting directly on the focal locus is more important than linked selection in determining introgression outcomes for individual loci.

Note that in the present example, the recipient population is sufficiently large or recombination between neighboring selected variants sufficiently frequent that individual variants can be isolated. More generally, selection and recombination are expected to weed out regions of some typical length *l*′ from within successful blocks, where *l*′ would depend on population size *N* and the growth rate of the successful block. The selective effects associated with replicable loci would then depend on the distribution of fitness effects of segments of length *l*′ rather than the DFE of individual variants. Understanding how *l*′ depends on population size *N* and *V*_0_ is non-trivial and we do not attempt to address this question here.

### Dependence of introgression probabilities on the size of the recipient population

The size of the recipient population is important in determining the probability of stochastic loss, and hence the extent to which the spread of an advantageous allele is predictable. The fixation probability of an unlinked allele with selective effect *s*, introduced into a population of size *N* at a frequency *p*_0_, was derived by Kimura (1957). This fixation probability approaches a limit that is independent of *N* for *N_s_*≫ 1: when a single copy of the allele is introduced (*p*_0_=1/2*N*), the asymptotic fixation probability is 2*s* for *s*≪1. Thus, most beneficial alleles fix in a single population. On the other hand, when a beneficial allele is introduced at a fixed frequency *p*_0_, its asymptotic fixation probability is 1 for large *N*. In this case, establishment of the beneficial allele is almost certain in the large *N* limit.

To what extent are these simple, single-locus predictions relevant to our model, where the introduced genome consists of multiple, linked, beneficial and deleterious loci? We investigate the dependence of long-run introgression probabilities of different single-locus variants on *N*, by comparing introgression of the same genome into populations of different sizes for two scenarios— one, where a single copy of the block is introduced into the population (*p*_0_=1/2*N*, see fig. 5A) and the second, where multiple copies of the same block are introduced at a fixed initial frequency (*p*_0_ independent of *N*, fig. 5B). Figures 5A and 5B show the introgression probabilities (at *t*=24000) of the single-locus variants versus their expected growth rates in the two cases, for various population sizes. Solid lines show the predictions of a modified version of Kimura’s formula, where the introgression probability *P_i_* of the introduced variant at locus *i* is assumed to be:

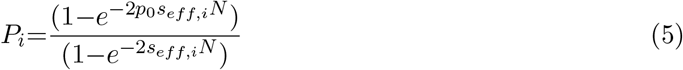

where *s_eff,i_* is the expected growth rate at locus *i*, rather than the selective effect of the locus.

**Figure 5:**
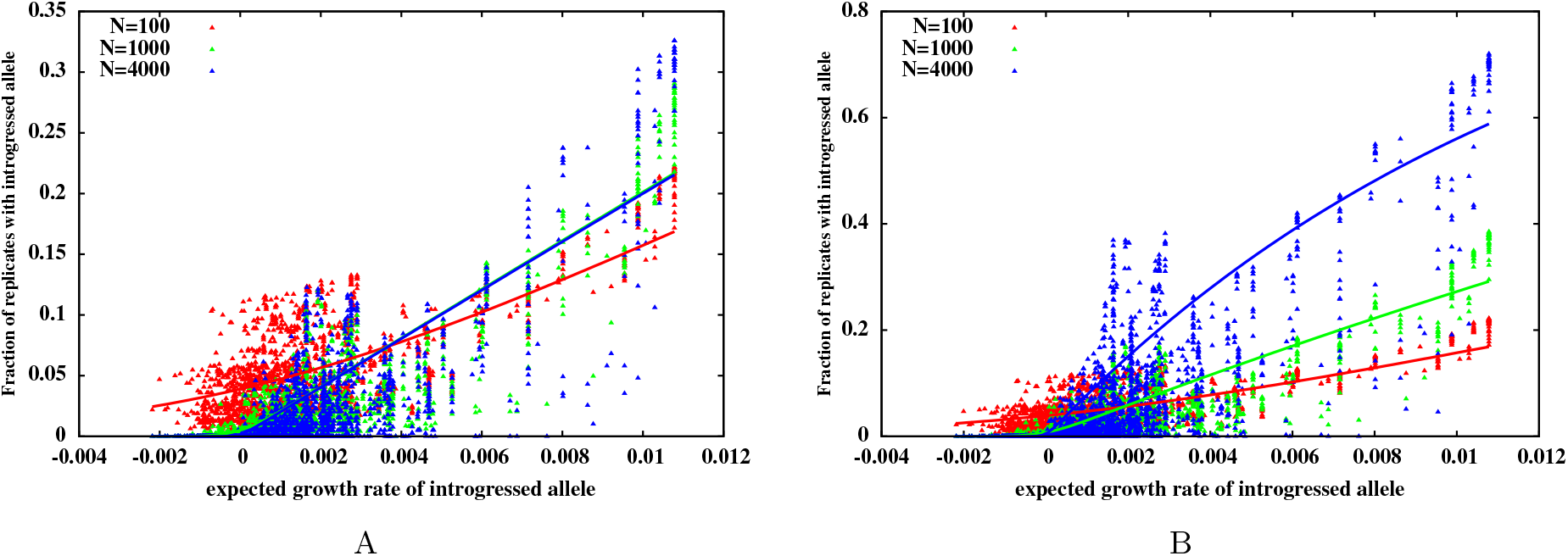
Introgression probabilities of single-locus variants versus their expected growth rates, for recipient populations of different sizes when (A) a single copy of the introduced block enters the population at *t*=0 (B) multiple (2*p*_0_*N* with *p*_0_ =0.005) copies of the block are introduced at *t*=0. Each point represents the introgression probability of a single-locus variant, as obtained from individual-based simulations for *N*=100 (red), *N*=1000 (green) and *N*=4000 (blue). Lines represent predictions of eq. (5). The same block (with *S*_1, *L*_=0, *y*_0_=0.4, 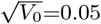, *L*=2048) is introduced in each simulation. Note the greater scatter in introgression probabilities (among loci with the same growth rate) for larger *N*.

Interestingly, the *average* variation of introgression probabilities as a function of the expected growth rate is captured quite well by eq. (5). This suggests that clusters of loci identified as having the same expected initial growth rate are, in some sense, natural linkage units into which the genome can be decomposed. To a first approximation, each such unit can then be considered as unlinked to others. Note, however, that there is considerable scatter of the introgression probabilities (about the average), especially in large populations. As we argue above, the differential introgression of tightly linked loci within a genomic fragment, depends on the time to fixation of the fragment relative to the time at which fine-scale recombination within the fragment becomes relevant, which in turn depends on population size *N*.

We now examine this dependence on *N* in more detail for the case of an introduced block with *L*=3 loci. Assume that the selective effects at the three loci satisfy *s*_1_>0, *s*_2_<0, *s*_3_>0, such that the expected growth rate of the full block is higher than the expected growth rates of any single or double locus fragments of the block. However, if recombination were to bring together introgressing variants at the first and third locus on a single block which lacks the deleterious (*s*_2_<0) variant in the middle, then this double recombinant would be fitter than the introduced parent block, and can supplant the latter.

An approximate expression for the fixation probability of this ‘101’ recombinant (carrying introgressed variants at the first and third loci) can be obtained using the semi-deterministic approach of Hartfield and Otto (2011) as follows (details in Appendix A). The 101 recombinant is mostly generated by recombination between between the full parent block, and a descendant block carrying just one introgressed variant (either at the first or the third locus). The frequency 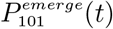 of such recombination events (that lead to the emergence of the 101 recombinant in generation *t*) can be computed by assuming deterministic dynamics for genotypic frequencies. We then use a branching process framework to calculate the probability *P_fix_*(*t*) that a 101 recombinant emerging at time *t*, fixes in the population in the long run. Note that the time at which the recombinant emerges determines its fitness relative to the recipient population, since the mean fitness of the recipient population is itself changing as the introgressed genome spreads through it.

The probability that none of the 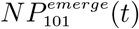 recombinants generated in generation *t* fix is then: 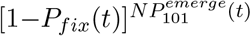, which is approximately 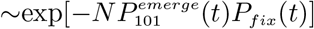. Further, if double recombinants emerge and establish rarely, then the net probability that no double recombinant fixes in a population of size *N* is just 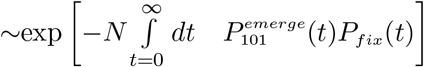. This implies that the double recombinant fixes with probability 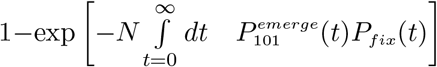 which increases with the size of the recipient population, and approaches 1 for large *N*, for a fixed *p*_0_. The fixation probability of the 101 recombinant in individual-based simulations matches the semi-deterministic prediction quite well (see Appendix A).

Thus, large populations typically have a larger number of recombinants that combine various fit fragments of a successful sub-block (before it fixes). This makes it increasingly probable that at least one of these recombinants establishes and supplants the parent sub-block in the population, thus eliminating deleterious variants embedded in the parent sub-block.

### Role of net selective effect of introduced block

So far we have considered the case where the introduced block is neutral with respect to the recipient population (*S*_1, *L*_=0). However, when the introduced block is deleterious as a whole (*S*_1, *L*_<0), any of its constituent fragments (even those which have a high, positive expected growth rate in the long run) experience an initial selective disadvantage. Many of the 2*p*_0_*N* copies of the introduced block (that carry this fragment) are lost from the population before the fragment can separated from the deleterious background and be amplified by selection, which suggests that the actual introgression probability in this case would be lower than that predicted by eq. (5).

Figure 6 compares introgression probabilities obtained from simulations with the predictions of eq. (5) for a neutral (fig. 6A), beneficial (fig. 6B), and deleterious (fig. 6C) block. As expected, eq. (5) broadly captures the variation of single-locus introgression probabilities along the neutral block, but systematically overestimates (or underestimates) these when the block is deleterious (or beneficial). Note that introgression probabilities of single-locus variants are significantly correlated with the expected growth rate, even for blocks with *S*_1, *L*_ ≠ 0 (where the correlation coefficient is *r*=0.77 for the beneficial block shown in fig. 6B and somewhat lower at *r*=0.51 for the deleterious block in fig. 6C). However, eq. 5 no longer accurately describes the relationship between the two quantities.

**Figure 6:**
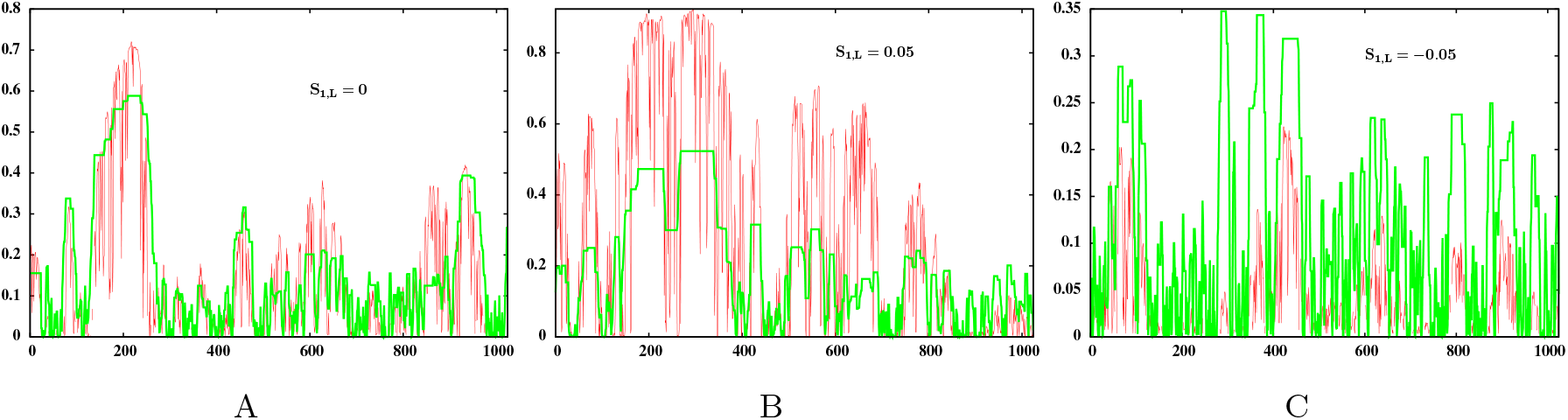
Introgression probability of the single locus variant at locus *i* on the introduced block, versus *i*, as obtained from individual-based simulations (red) and as predicted by eq. (5) (green), for an introduced block that is (A) neutral with respect to the recipient population (*S*_1, *L*_=0) (B) beneficial (*S*_1, *L*_=0.05) (C) deleterious (*S*_1, *L*_=−0.05). Each of the three blocks has map length *y*_0_=0.4 and *L*=2048 loci with selective effects drawn from a distribution with mean *μ*=*S*_1, *L*_/*L* and variance *σ*^2^=0.001/*L* (subject to the constraint that the effects of all loci add up to *S*_1, *L*_. In any set of simulations, identical copies of the block are introduced at frequency *p*_0_=0.005 in a diploid population of size *N*=4000. For ease of visualization, only half of the genome is shown. Note that eq. (5) broadly captures variation of introgression probability along the introduced block for the *S*_1, *L*_=0 block, but significantly underestimates or overestimates probabilities when *S*_1, *L*_≠0.

More generally, since the rate of increase of a sub-block itself changes over time, before approaching a constant (asymptotic) value (see fig. 1B), a more accurate predictive approach needs to explicitly consider the establishment of any fragment under a time-varying effective selection coefficient (see e.g., Uecker and Hermisson 2011) instead of a constant *s_eff_* (as used in eq. (5)). An interesting question is whether the effect of total fitness *S*_1, *L*_ of the introduced block can be partially captured by a constant factor (corresponding to a genome-wide barrier to gene flow), that attenuates (or inflates) the introgression probabilities of all fragments of the block by the same amount. Explicit expressions for barrier strength have been derived by Bengtsson (1985) and Barton and Bengtsson (1986) for the simpler case of a neutral locus embedded in a genome with all deleterious loci.

### Dependence of long-term selection response on population size and linkage

Directional selection on standing genetic variation results in a net phenotypic advance, which depends on population size *N*. In the case of selection on a very large number of polymorphic *unlinked* loci (as in the standard infinitesimal model), additive genetic variance declines at a rate ~1/2*N* per generation due to drift. This constrains the net advance, which scales *linearly* with *N* (Robertson 1960). The situation is more complex for selection on linked loci (Robertson 1970, 1977). Tightly linked variants would fix together rapidly in a small population, but be broken apart and seen by selection in a larger population. Thus, in this case, increasing population size tunes drift as well as the extent of hitchhiking. How does the resultant net advance under selection scale with population size in this model, for short versus long introduced blocks?

We investigate this by simulating a particular set of *L*=2048 loci, with net trait value *S*_1_=0, spread across a block of different map lengths *y*_0_. Multiple (i.e., 2*p*_0_*N*) copies of the block are introduced into a population of size *N* at *t*=0. We then measure the net advance under selection (defined as the average trait value in the limit *t*→∞) in a population, and average over replicate populations. To assess the effect of linkage on net advance, we do simulations where the same *L*=2048 loci are uniformly spaced across blocks with different *y*_0_; the allelic effects of the loci are scaled with 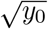 such that the genic variance per unit map length (*V*_0_=*σ*^2^*L*/*y*_0_) and the trait value relative to the total genic variance 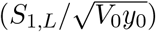 is the same for blocks of different map lengths. Figure 7 shows the average net advance versus population size *N*, for blocks of different lengths. The net advance is normalized by the probability of survival of at least some part of the introduced block.

**Figure 7:**
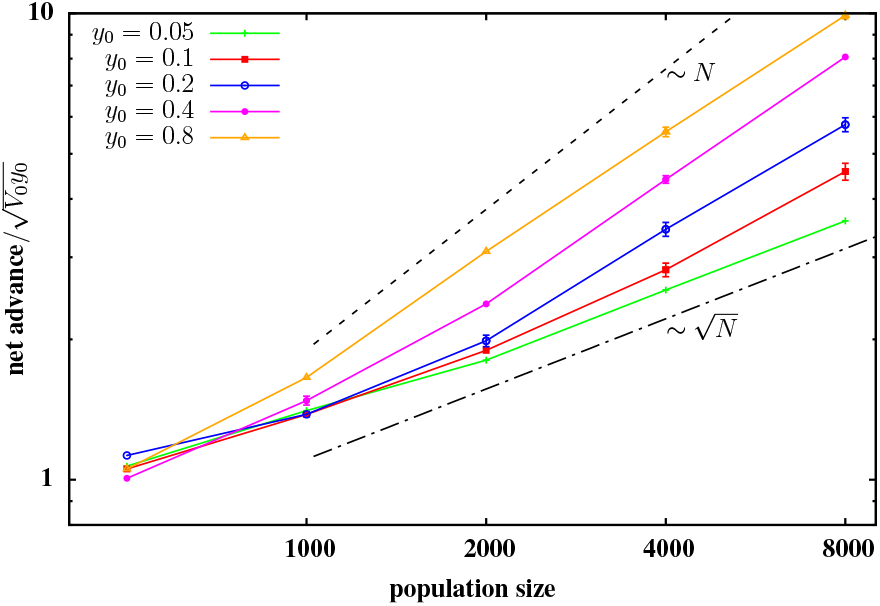
The net advance relative to 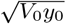, versus size *N* of the recipient population, for introduced blocks of different lengths *y*_0_. A given block consists of *L* uniformly spaced loci; the allelic effects are drawn from a Gaussian distribution with mean 0 and variance *σ*^2^=0.4/*L* (for the block of length *y*_0_=0.05), such that the effects across all loci sum to 0. For the other (longer) blocks, we scale the allelic effects with 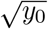, so that the genic variance per unit map length is the same for blocks of different map lengths. All points are obtained from individual-based simulations. The net advance scales sub-linearly with *N* for small *N*, but appears to approach a linear dependence for larger *N*; the approach to ~*N* behaviour is faster for longer blocks.

Figure 7 shows that the net advance scales sub-linearly with *N* for the shortest blocks (tightly linked loci), but approaches a linear dependence with increasing block length. Based on the above arguments, we expect efficient fine-scale separation of beneficial and deleterious loci to also occur within short blocks in sufficiently large populations, resulting in an asymptotically linear scaling of net advance with *N*. However, this is difficult to confirm in simulations. More importantly, this limit might not be realized in typical scenarios involving populations of a few thousand (as shown in fig. 7), when there is tight linkage between introduced variants; thus, in practice, one would often observe a sub-linear scaling of net advance with the size of the recipient population.

## Discussion

### Selection on linkage blocks

The nature of genetic change (substitutions versus minor shifts in allele frequency) at individual loci during polygenic adaptation and the resultant genomic signatures of such adaptation have attracted much interest recently (Pritchard et al 2010). A key challenge is to disentangle the effect of selection at a given locus on the allele frequency dynamics of that locus, from the effect of linked selection due to nearby loci. Our analysis shows that these two kinds of selection act over different time scales, and together shape the fixation probability of different variants on a single genome introduced into a genetically homogeneous population. We identify sets of loci that share the same short-term fate, by decomposing the genome into contiguous blocks that grow faster (under directional selection) than any parent block containing them. Each such block acts as a linkage unit: the initial introgression of loci within a block is governed by the rate of spread of the block (rather than the selective effects of individual loci), and is largely independent of other such blocks.

While the expected growth rates of linkage blocks provide a useful first estimate for the variation of introgression probability along the genome, a more refined estimate needs to account for several other factors such as the fine scale variation within successful fragments, the effects of linkage between multiple successful fragments, and the effect of the background genome, which could attenuate or inflate the effective number of blocks introduced into the population.

The observation that a recombining genome under selection can be viewed as a collection of unlinked, effectively asexual segments or ‘linkage blocks’ is quite powerful, and allows for the calculation of neutral diversity within these segments using theoretical results for asexual populations (Neher et al 2013; Weissman and Hallatschek 2014; Good et al. 2014). Note that in our formulation, the intrinsic growth rate of a fragment of map length *y* and selective effect *S_y_* is proportional to *S_y_*−*y*, which is of the order of 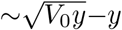, if the full introduced block is neutral (*S*_1, *L*_=0). This is maximum for a characteristic map length *y*\*∝*V*_0_ or alternatively, for a characteristic number of loci given by *L*\*~*σ*^2^/*c*^2^, where *σ* is the variance of fitness effects of individual variants and c the rate of recombination between adjacent selected variants. Thus, on average, blocks of length *y*\* should grow faster than any parent block containing them. This heuristic derivation for the characteristic linkage scale is consistent with the expectations of Neher et al (2013) (see eq. 5 in that study), modulo a logarithmic correction arising from the dependence of fitness variance on population size.

### Selection on individual loci and net advance under selection

While the expected growth rate of linkage blocks show a high correlation with the long-term fixation probability of individual variants within these (fig. 3B), and can capture the coarse-grained variation of introgression probabilities along the introduced block reasonably well (see fig. 6A), there can be considerable fine-scale variation of introgression probabilities of individual variants within each such linkage unit. In fact, this fine-scale variation often reflects differences in selective effect of variants *within* a successful block (fig. 3D). As we illustrate by means of a toy example, the efficacy of selection at fine scales depends on the rate at which a fit block fixes in the population, relative to the rate at which recombination can generate and bring together fragments of the block into even fitter combinations that lack some of the deleterious variants present in the original block. In larger populations, the latter rate is larger, resulting in finer scale sifting of deleterious variants with increasing population size at long time scales.

The extent to which selection can eliminate individual deleterious variants embedded within a favourable background determines the net response under selection. Robertson (1970) derived approximate expressions for the net advance due to selection acting on standing variation contributed by linked loci of equal effects (spread over map length *l*) and found that the net advance in a population of size *N* was significantly reduced relative to the advance under free recombination, while *l* is less than the selection intensity. Importantly, for tight linkage, net advance scales sub-linearly with *N* for small *N*, and approaches a linear dependence for larger *N*. Our results are broadly consistent— the net advance increases with increasing map length (fig. 7). Further, the net advance scales approximately linearly with *N* for the longest block (weakest linkage), but shows a much weaker (sub-linear) dependence on *N* for tighter linkage between loci (see fig. 7).

### Selective effects associated with highly replicable loci

Our analysis shows that even highly replicable substitutions may be associated with a wide range of selective effects and can even be deleterious, especially if the density of selected variants on the genome is high (see fig. 3C or fig. 4). The fraction of deleterious substitutions during adaptation is lower when the trait value is determined by fewer, more widely-spaced loci of stronger effects. In this case, a typical linkage block contains fewer (∝*σ*^2^/*c*^2^) loci, which implies that the selective effect of a locus contributes more to and is thus more strongly correlated with its expected growth rate. Thus deleterious loci are less likely to be present in an initially successful parent block and hitchhike to high frequencies during the initial phase of introgression. Moreover, even those deleterious variants that attain high frequencies have a higher chance of being eliminated at longer time scales once fine scale recombination starts isolating individual loci, due to stronger selection against individual variants. Note that a wider DFE actually results in stronger hitchhiking and a higher fraction of deleterious substitutions unless the density of selected variants is correspondingly low (fig. 4).

Interestingly, differences in replicability among *tightly linked* individual variants are informative about the effect sizes of these variants— missing variants or polymorphic variants (that are declining in frequency) within genomic regions with high overall introgression probability are typically associated with deleterious effects. By contrast, differences in introgression probabilities of loosely linked loci (such as those belonging to different linkage blocks) are more informative about the variation in the strength of linked slection along the genome and cannot be explained by differences in selective effects of individual loci (contrast figs. 3D and 3C). In general, our study underlines the need for a cautious approach towards interpreting patterns of introgression or adaptation in replicates, and suggests that it is more difficult to ascribe adaptive significance to individual substitutions than to clusters of such substitutions. While genomic regions with multiple replicated substitutions (which correspond to successful blocks in our model) are typically important for adaptation, individual substitutions within these may be nearly neutral (even if they occur across replicates, see fig. 3D).

### Interpreting genomic islands of divergence

Genomic regions where the native haplotype persists (i.e., for which the introduced haplotypes fail to introgress) would appear as islands of divergence between the recipient and source population (Strasburg et al 2012). In our model, such regions are associated with introduced fragments that have a *negative* expected growth rate in the recipient population. Note that such fragments need not have deleterious effects; negative growth rates can also be associated with moderately beneficial fragments that are nevertheless broken up faster by recombination than they are amplified by selection. Such fragments may be completely lost from the population, especially when introduced at a low frequency into the recipient population. Thus, in this model at least, the hybridization history (steady gene flow versus sporadic migration events) can be an important factor determining whether extended regions of differentiation are necessarily correlated with barriers to gene flow.

### Replicability during adaptation to a phenotypic optimum

Our model considers an additive trait under directional selection and thus ignores epistasis and genetic redundancy, which can strongly influence replicability (Yeaman et al. 2015). A common scenario in which epistatic constraints influence adaptation is one where the additive trait is under stabilizing selection towards a different optimum in the recipient population. In this case, if introduced haplotypes are far from the selection optimum, then they effectively experience directional selection, causing adaptive fragments within these to be amplified in much the same way as in the present model. In the long run, closer to the optimum, the redundancy of different genomic fragments that make similar contributions to the phenotype may play an important role. Understanding how the initial spread of adaptive fragments constrains long-term introgression patterns along the genome in this case is an interesting direction for future work.

### Genetically heterogeneous donor and recipient populations

Our model assumes that the recipient population is genetically homogeneous; further, even the initial hybridization event involves introduction of multiple copies of the *same* genomic block into the recipient population. These assumptions thus effectively ignore variation within the source and recipient population, and just focus on fixed differences between the two. Most theoretical work on introgression and barriers to gene flow makes similar assumptions (Barton and Bengtsson 1986; Uecker et al. 2015). However, genetic heterogeneity within the donor and recipient populations is likely to qualitatively impact patterns of introgression and the extent to which these are convergent across replicate populations, and is thus a natural extension to consider.

The present analysis provides some intuition in a scenario where a few non-identical haplotypes are introduced into a genetically homogeneous recipient population: genomic fragments that have high expected growth rates and are present in multiple copies among the introduced haplotypes are expected to introgress with high probability. However, genetic variation within the donor and recipient populations can have other important consequences. Small recipient populations may exhibit significant inbreeding depression due to segregation of deleterious recessive alleles. Alleles from a diverged population then have a heterotic advantage even if they are deleterious within the recipient population; in fact, heterosis has been suggested as a possible reason for the persistence of Neanderthal-derived variants in modern human populations (Harris and Nielsen 2016).

### Efficacy of selection

We have considered the contribution that a single introduced genome makes to a homogeneous population. A natural extension is to consider the efficacy of selection within a heterogeneous population. Initially, there is a set of genomes, each carrying alleles at very many loci that have a distribution of effects, both positive and negative. We can focus on one of these genomes, and ask what contribution it makes to the final selection response; the expected response of the whole population is the sum of the expected contributions of each initial genome. The key differences from the analysis presented here are that a genome finds itself competing against other genomes that are themselves improving under selection, and that it finds itself associated with other genomes that have random effects on the trait. A first step might be to simply treat the rest of the population as having a mean and variance determined by classical quantitative genetics, and ask whether an analysis focused on a single genome in this variable background can give a good approximation.

We have contrasted two regimes: one where the fate of an allele depends on its own selective effect, and another, where selection acts on fragments of genome whose effects on fitness depend on very many loci. This contrast is closely related to the more practical question of when we can detect individual causal alleles, by genetic mapping or an association study, or instead, can only attribute genetic variance to regions of genome. An important question for the future is to find ways to determine when it is appropriate to deal with discrete loci, rather than take a statistical approach: or better, how to combine these two representations of genetic variation in order to understand real genetic architectures.

## Appendix

### A. Introgression of a block with 3 loci: semi-deterministic approximations

Consider a genotype ‘111’ introduced at frequency *p*_0_ in a population of *N* individuals, all having genotype ‘000’. For simplicity, we consider haploid individuals, but the analysis can be generalized to the case with diploids. In the following, ‘1’ will always denote the introduced allele and ‘0’ the native allele. The native alleles at each of the three loci are associated with zero selective effect. The introduced alleles at the three loci have selective effects *s*_1_, *s*_2_ and *s*_3_; the net selective effect of the introduced genotype is *s*=*s*_1_+*s*_2_+*s*_3_. The introduced genotype is of the type +−+, i.e., with *s*_1_>0, *s*_2_<0 and *s*_3_>0. We now derive an approximate expression for the probability of fixation of the double recombinant ‘101’ (i.e., +0+), which is the fittest possible recombinant, using the semi-deterministic approach employed by Hartfield and Otto (2011).

We consider frequencies of genotypes generated by a single recombination event. These single recombinants are: 100, 110, 001, 011. The genotypes 110 and 011 will typically be quite rare, as they have lower fitness than the genotypes 001 and 100, as well as the introduced genotype 111. As a first approximation, we assume that their frequency is zero. Then we need only track three genotypes: 001, 100 and 111 (along with the native genotype 000), before the genotype 101 emerges. Under deterministic dynamics, and assuming *s,c*≪1, such that all second order terms in *s* and *c* can be neglected, the frequencies of these genotypes evolve as:

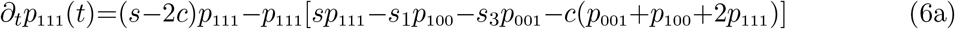

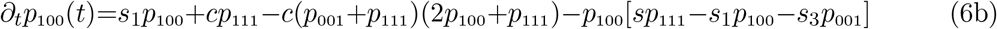

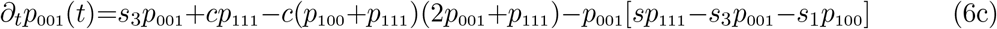

These equations can be solved numerically to obtain various genotypic frequencies as a function of time.

The fraction of mating events that generate the recombinant 101 in generation *t* is *cp*_111_(*t*)[*p*_100_(*t*) + *p*_001_(*t*)]+2*cp*_100_(*t*)*p*_001_(*t*). Since the frequencies *p*_001_(*t*) and *p*_100_(*t*) are typically quite small, we can approximate the probability of emergence 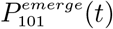 of the recombinant 101 by 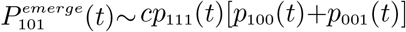. Note that 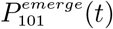 is the probability that a mating event in generation *t* results in a 101 recombinant; it is not the frequency of 101 genotypes in generation *t*.

We can now calculate the fixation probability *P_fix_*(*t*) of a 101 recombinant generated in generation *t*, by approximating its spread as a time-inhomogeneous branching process, where the relative selective advantage of this recombinant changes over time, due to the changing mean population fitness (Hartfield and Otto 2011). Then *P_fix_*(*t*) follows (see Barton 1995):

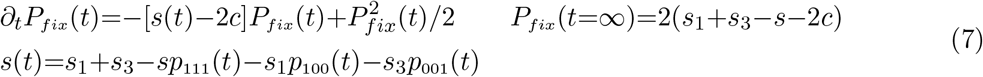

Equation (7) can be solved numerically to obtain *P_fix_*(*t*). Then the probability that none of the 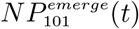 recombinants (that emerge in generation *t*) fix is just: 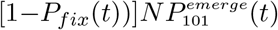, which can be approximated by 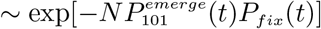. Further, the net probability that no double recombinant fixes in a population of size *N* is 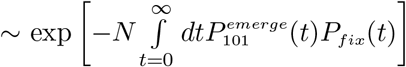. The key assumption is that instances of double recombinants emerging and establishing are rare, so that all such instances can be treated as independent.

We compare predictions of this semi-deterministic analysis with results of individual-based simulations of populations of different sizes (fig. 8), and find that there is good agreement between the two. The analysis above assumes deterministic dynamics of the introduced block and other genotypes produced by single crossover events. A more careful analysis can be done to account for stochastic effects in the initial phase, when the introduced block and its descendants are rare and may be lost by chance.

**Figure 8:**
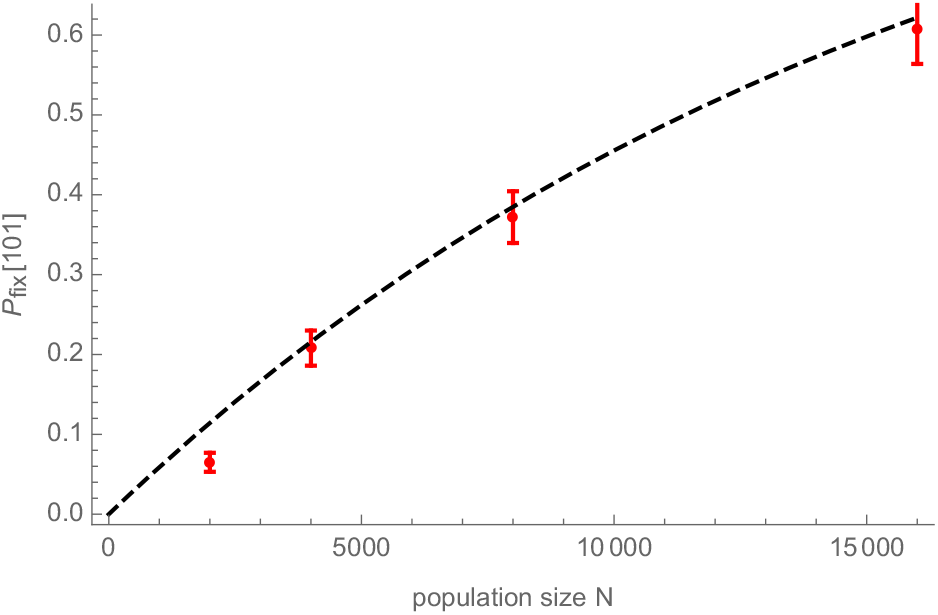
Probability that the ‘101’ recombinant fixes in a population with *N* individuals, versus *N*. The results of individual-based simulations (points) show good agreement with the prediction of the semi-deterministic analysis described above (dashed line). The introduced block has 3 loci with *s*_1_=0.0012, *s*_2_=−0.0006 and *s*_3_=0.0014; the rate of recombination between adjacent loci is *c*=0.0002; the initial frequency of the introduced genotype is *p*_0_ =0.006. The fixation probability is normalized with respect to the probability that at least some part of the introduced block survives.

